# Speed tuning to real-world-and retinal motion in cortical motion regions

**DOI:** 10.1101/449256

**Authors:** Didem Korkmaz Hacialihafiz, Andreas Bartels

## Abstract

Motion signals can arise for two reasons in the retina: due to self-motion or due to real motion in the environment. Prior studies on speed tuning always measured joint responses to real and retinal motion, and for some of the more recently identified human motion processing regions, speed tuning has not been examined in at all. We localized motion regions V3A, V6, V5/MT, MST and cingulate sulcus visual area (CSv) in 20 human participants, and then measured their responses to motion velocities from 1-24 degrees per second. Importantly, we used a pursuit paradigm that allowed us to quantify responses to objective and retinal motion separately. In order to provide optimal stimulation, we used stimuli with natural image statistics derived from Fourier scrambles of natural images. The results show that all regions increased responses with higher speeds for both, retinal and objective motion. V3A stood out in that it was the only region whose slope of the speed-response function for objective motion was higher than that for retinal motion. V6, V5/MT, MST and CSv did not differ in objective and retinal speed slopes, even though V5/MT and MST tended to respond more to objective motion at all speeds. These results reveal highly similar speed tuning functions for early and high-level motion regions, and support the view that human V3A encodes primarily objective rather than retinal motion signals.

## Introduction

Motion perception is a crucial function of the visual cortex. Perceiving the speed of motion is important for many everyday tasks of the visual system, such as pursuing a target or interacting with a moving object, or even for compensating self-induced retinal motion with non-retinal signals. Although visual motion perception and processing have been widely studied, our understanding of speed processing is still limited. Most prior studies on speed processing focused on V5/MT both in humans and primates. Majority of V5/MT neurons were shown to be speed selective ((<u>Maunsell & Van Essen, 1983b</u>; <u>Perrone & Thiele, 2001</u>) but also see (<u>Priebe, Cassanello, & Lisberger, 2003</u>)) and V5/MT plays an important role in speed perception and discrimination, which was shown by lesion studies (<u>Dursteler & Wurtz, 1988</u>; <u>Newsome, Wurtz, Dürsteler, & Mikami, 1985</u>; <u>Orban, Saunders, & Vandenbussche, 1995</u>; <u>Pasternak & Merigan, 1994</u>; <u>Yamasaki & Wurtz, 1991</u>). Neurons are clustered according to preferred speed with no columnar organisation for speed in V5/MT (<u>Liu & Newsome, 2003</u>). The range of preferred speeds of neurons were reported differently in different studies, such as between 2-256 deg/s (<u>Maunsell & Van Essen, 1983b</u>) or between 5-150 deg/s (<u>Rodman & Albright, 1987</u>). The peak of preferred speeds across neurons was reported as 32 deg/s (<u>Maunsell & Van Essen, 1983b</u>) and speed representation in V5/MT is thought to be in logarithmic scale (<u>Nover, Anderson, & DeAngelis, 2005</u>). Speed perception is shown to be consistent with average responses from V5/MT neurons of macaques (<u>M. M. Churchland & Lisberger, 2001</u>; <u>Priebe & Lisberger, 2004</u>), although it has also been shown in monkeys that overall sensitivity of V5/MT neurons to speed discrimination is less than the behavioural responses (<u>Liu & Newsome, 2005</u>). Despite the vast number of studies to characterize speed responses of V5/MT neurons in non-human primates, our knowledge about the speed responses in human V5/MT is limited.

Previous human psychophysics studies on speed discrimination show that humans can discriminate even small differences in speed when the speed is between 4-32 deg/s (<u>Beauchamp, Cox, & DeYoe, 1997</u>; <u>Orban, de Wolf, & Maes, 1984</u>). Speed coding in human V5/MT using an adaptation paradigm was shown by fMRI findings (<u>Lingnau, Wall, & Smith, 2009</u>) and application of TMS (transcranial magnetic stimulation) over human V5/MT area induced deficits in speed perception, revealing an important role of V5/MT in speed perception in humans (<u>McKeefry, Burton, Vakrou, Barrett, & Morland, 2008</u>). Using fMRI, Chawla and colleagues found that optimal speed responses in human V5/MT were between 7 deg/s and 30 deg/s in one study (<u>Chawla, Phillips, Buechel, Edwards, & Friston, 1998</u>) and between 4 deg/s and 8 deg/s in another study (<u>Chawla et al., 1999</u>). Additionally, some researchers investigated how other image features effect the speed perception. For instance, in both humans and monkeys, it has been shown that changes in contrast result in alterations in both perception of speed and speed tuning, such that low contrast causes lower speed perception and shifts in neural responses to lower speed tuning and firing rate (<u>Krekelberg, Boynton, & van Wezel, 2006</u>).

Previous studies also examined speed-dependent responses in another motion responsive region, V3A (<u>Arnoldussen, Goossens, & van den Berg, 2011</u>; <u>Chawla et al., 1999; Chawla et al., 1998</u>; <u>McKeefry et al., 2008</u>). In macaques, almost all V3A neurons are reported to have speed sensitivity, and this sensitivity is present for a wide range of speeds, even faster than 50 deg/s (<u>Galletti, Battaglini, & Fattori, 1990</u>). In humans, fMRI results showed that V3A has speed responses similar to MT/V5, although in V3A, optimal speed range is reported to be between 4-16 deg/s (<u>Chawla et al., 1999; Chawla et al., 1998</u>) and application of TMS over V3A has been shown to impair speed perception (<u>McKeefry et al., 2008</u>). Particularly, V3A is responsive to headcentric speed of motion (<u>Arnoldussen et al., 2011</u>). Apart from its speed tuning, human V3A is just as sensitive to motion as V5+/MT+ (<u>Bartels, Zeki, & Logothetis, 2008</u>; <u>Tootell et al., 1997</u>). Importantly though, it differs substantially from the human V5/MT+ complex in that it is primarily driven by real-world motion, with nearly no response to retinal motion per se (<u>Fischer, Bulthoff, Logothetis, & Bartels, 2012a</u>). Consistent with this, in monkey, it contains a large fraction of real-motion neurons (<u>Galletti & Battaglini, 1989</u>; <u>Galletti et al., 1990</u>; <u>Ilg, Schumann, & Thier, 2004</u>).

All prior studies examining speed responses measured joint responses to objective and retinal motion, since participants fixated while speed of the (objective) background motion varied. This induced motion on the retina that was the result of physical motion on the screen. However, during eye-and head-movements, the visual system continuously compares retinal motion with efference copies of the eye, in order to estimate world-centered motion. This allows it to create a stable perception of the world despite retinal motion, or to infer object-motion also when eye-movements reduce retinal motion to zero. Previously, human visual areas V3A and V6 were shown to compensate for self-induced retinal motion and encode almost exclusively objective motion during smooth pursuit eye movements (<u>Fischer et al., 2012a</u>). To a weaker extent, also CSv was shown to discount self-induced retinal motion (<u>Fischer, Bulthoff, Logothetis, & Bartels, 2012b</u>). CSv responds to egomotion related visual signals (<u>Cardin & Smith, 2010</u>; <u>Fischer et al., 2012b</u>; <u>Wall & Smith, 2008</u>), heading direction (<u>Furlan, Wann, & Smith, 2014</u>), and vestibular signals (<u>Smith, Wall, & Thilo, 2012</u>).

V6 is located in parieto-occipital sulcus (POS) (<u>Pitzalis et al., 2006</u>), shows retinotopic organisation both in humans and macaque (<u>Galletti, Fattori, Gamberini, & Kutz, 1999</u>; <u>Pitzalis et al., 2006</u>; <u>Pitzalis et al., 2010</u>) with a stronger representation of periphery, although fovea is also represented (<u>Pitzalis et al., 2006</u>) and responds preferentially to wide field stimulation, and responds to ego-motion compatible coherent motion (<u>Cardin & Smith, 2010</u>; <u>Pitzalis et al., 2006</u>; <u>Pitzalis et al., 2013</u>; <u>Pitzalis et al., 2010</u>). V6, together with V3A, distinguishes real motion from self-induced retinal motion (<u>Arnoldussen et al., 2011</u>; <u>Fischer et al., 2012a</u>). Moreover, V6 is involved in processing of head-centric translation and rotation speed (<u>Arnoldussen et al., 2011</u>; <u>Arnoldussen, Goossens, & van den Berg, 2015</u>). For none of these regions speed responses have been measured separately for retinal and objective motion.

Here we used a pursuit paradigm that allowed us to do just that (<u>Fischer et al., 2012a</u>): participants fixated a disc that was either stationary or moved along a circular trajectory around the screen. At the same time, either a stationary background was shown or one that moved on the same circular trajectory. This 2 x 2 factorial design led to four conditions that allowed separating responses to retinal or objective motion in the absence of pursuit-related confounds. We applied this paradigm at six different levels of speed (1, 2, 4, 8, 16, and 24 degrees per second) in order to obtain speed-tuning profiles for retinal and objective motion for separately localized visual motion regions V5/MT, MST, V3A, V6 and CSv. The background stimulus was derived from Fourier scrambles of natural images in order to match natural image statistics. This provides a closer match to the spatio-temporal exposure in real-life and has been shown to provide more optimal stimuli for the visual system (<u>Kayser, Kording, & Konig, 2004</u>; <u>Vinje & Gallant, 2000</u>). Further, speed responses in MT were enhanced when the stimuli had multiple spatial frequencies and this was thought to be a result of natural scenes containing multiple spatial frequencies (<u>Priebe et al., 2003</u>; <u>Priebe, Lisberger, & Movshon, 2006</u>). We found that only in V3A, the slope of objective motion speed was significantly higher than the slope of retinal motion. Additionally, all regions were modulated by the speed of both types of motion.

## Materials and Methods

### Participants

20 healthy participants with normal or corrected vision (14 female, 7 male, 4 left handed, age between 18 and 37 (average: 26), 1 author) took part in this study. All participants gave written informed consent before the experiment and were compensated for their participation. The study was approved by the local ethics committee of the University Hospital of Tübingen.

### Main Experiment

#### Stimuli and Paradigm

We used a 2 x 2 factorial pursuit paradigm that allowed us to measure responses to objective and retinal motion separately (<u>Fischer et al., 2012a</u>). This paradigm was applied using 6 different levels of speed (1, 2, 4, 8, 16, and 24 degrees per second). The factors pursuit (on/off) and objective planar motion (on/off) resulted in the following 4 conditions: fixation on a static background, fixation on a moving background, pursuit on a static background, and pursuit on a moving background. In the last condition, the pursuit trajectory was locked to that of the background, thus nulling retinal motion.

The motion trajectory of the background and pursuit followed a circular path with a radius of 4 degrees (1/4^th^ of the screen height, with screen dimensions of 22 x 16 degrees). The motion radius was chosen such that the area of controlled visual stimulation was maximal: the nearest border to the screen edge was at all times further away than 4 visual degrees, leading to controlled visual stimulation within at least 8 x 8 visual degrees (<u>Fischer et al., 2012a</u>). The rotation direction and starting point of the fixation cross was randomized and counter balanced within each participant. Note that the upper speed limit of 24 deg/s was set following piloting by the authors showing that this speed could still be reliably pursued on its circular trajectory, while considerably higher speeds could not.

For the background, 100 pink noise images were created from 100 natural scene images using Fourier phase scrambling. The images were converted to greyscale and contrast and luminance were matched across the images before phase scrambling. Stimuli were chosen randomly for each trial and back-projected with 1024 x 768 pixels resolution and 120 Hz refresh rate. The size of the images was 2048 x 1536 pixels, with 737.5 cd/m2 luminance and 147.5 root-mean-square (RMS) contrast.

#### Procedure

The experiment was presented in a block design manner; each condition lasted 12 s, and was preceded by 1 s fixation and followed by 2 s fixation, leading to an inter-trial interval of 15 s. The fixations at the beginning and end of each trials were presented in order to minimize confounds related to eye movements at the beginning of each trial since the location of starting point in each trial was randomized. Each condition was shown 4 times per run, yielding a total number of 17 blocks including one initial additional block for counterbalancing. We used back-matched pseudorandom sequences (<u>Brooks, 2012</u>) such that each condition was preceded equally often by all conditions. The initial block inserted for counterbalancing was discarded from further analyses. Each run started with 1.74 s grey screen with fixation and ended with 10 s of grey screen with fixation. There was a grey fixation disk (width: 0.74 degrees, luminance: 1153.7 cd/m2) with a fixation task (see below) at all times. The order of runs was also pseudorandomized for half of the participants and the flipped version of this run sequence was used for the other half.

In total, six runs were acquired from each participant, each run containing one speed level. The sequence of speed levels was random and counterbalanced across participants.

The main experiment was presented using Psychtoolbox 3.0 (<u>Brainard, 1997</u>; <u>Kleiner et al., 2007</u>) and Matlab 7.10.0 (<u>MATLAB, 2010</u>).

### Motion Localizer

In order to localize V5/MT, MST, CSv, V3A and V6, we used an independent motion localizer that was previously described (<u>Fischer et al., 2012a, 2012b</u>). It consisted of 7 conditions presented 12 s each in 7 counterbalanced repetitions: 3D full field motion (coherent expanding/contracting motion), random motion (with trajectories matched to 3D motion), right and left hemifield 3D full field motion (left or right 2/5^th^ of the screen), 2D planar motion with synced pursuit, static background with pursuit, and static baseline. In each condition except baseline, random patterns of black and white dots on a grey background were used as stimuli. The baseline consisted of blank grey screen. As in the main experiment, there was a grey fixation disk (width: 0.74 degrees) with the fixation task (see below). The stimuli were presented using Cogent 2000 developed by the Cogent 2000 team at the FIL and the ICN and Cogent Graphics developed by John Romaya at the LON at the Wellcome Department of Imaging Neuroscience (http://www.vislab.ucl.ac.uk/cogent_graphics.php) and Matlab 7.10.0 (<u>MATLAB, 2010</u>).

### Region of interest (ROI) definitions

ROIs were defined using the MarsBar toolbox as follows: MST was defined as ipsilateral response within the V5+/MT+ complex using hemifield motion versus baseline contrast, V5/MT was defined as contralateral response during the same contrast excluding MST voxels. V6 and CSv were localized using 3D coherent motion versus random motion. V3A was localized using 2D lateral motion with pursuit versus smooth pursuit with static background. For each participant, each region was localized using an individual p-value, and when ROIs could not reliably be detected it was not defined at all (<u>Fox, Iaria, & Barton, 2009</u>; <u>Murray & Wojciulik, 2004</u>).

### Fixation Task

In order to balance attention across conditions, participants were required to do a 1-back character-matching task, which was a randomly presented sequence of alphabetical characters displayed one at a time on the fixation disk. There was a repeating character between every 3 to 8 character presentations, which were reported by participants via button press. The timing of these button presses was included in the GLM analysis as a regressor.

### Image Acquisition

Data were acquired with a Siemens Magnetom PRISMA 3 Tesla scanner using a 64-channel phased-array head coil (Siemens, Erlangen, Germany). A gradient echo sequence consisting of T2* weighted images with the following parameters was used for functional scans: TR = 0.87 s, TE = 30 ms, flip angle = 57°, Generalized Autocalibrating Partially Parallel Acquisitions (GRAPPA) g-factor = 2, multi-band factor = 4, voxel size = 2 x 2 x 2 mm^3^. 56 slices were acquired in an interleaved order. T1 weighted anatomical images were acquired with a resolution of 1 x 1 x 1 mm^3^. The first 2 functional volumes of each run were discarded for T1 equilibration. Field map images were also acquired in order to correct for B_0_ field distortions.

**Figure 1.**
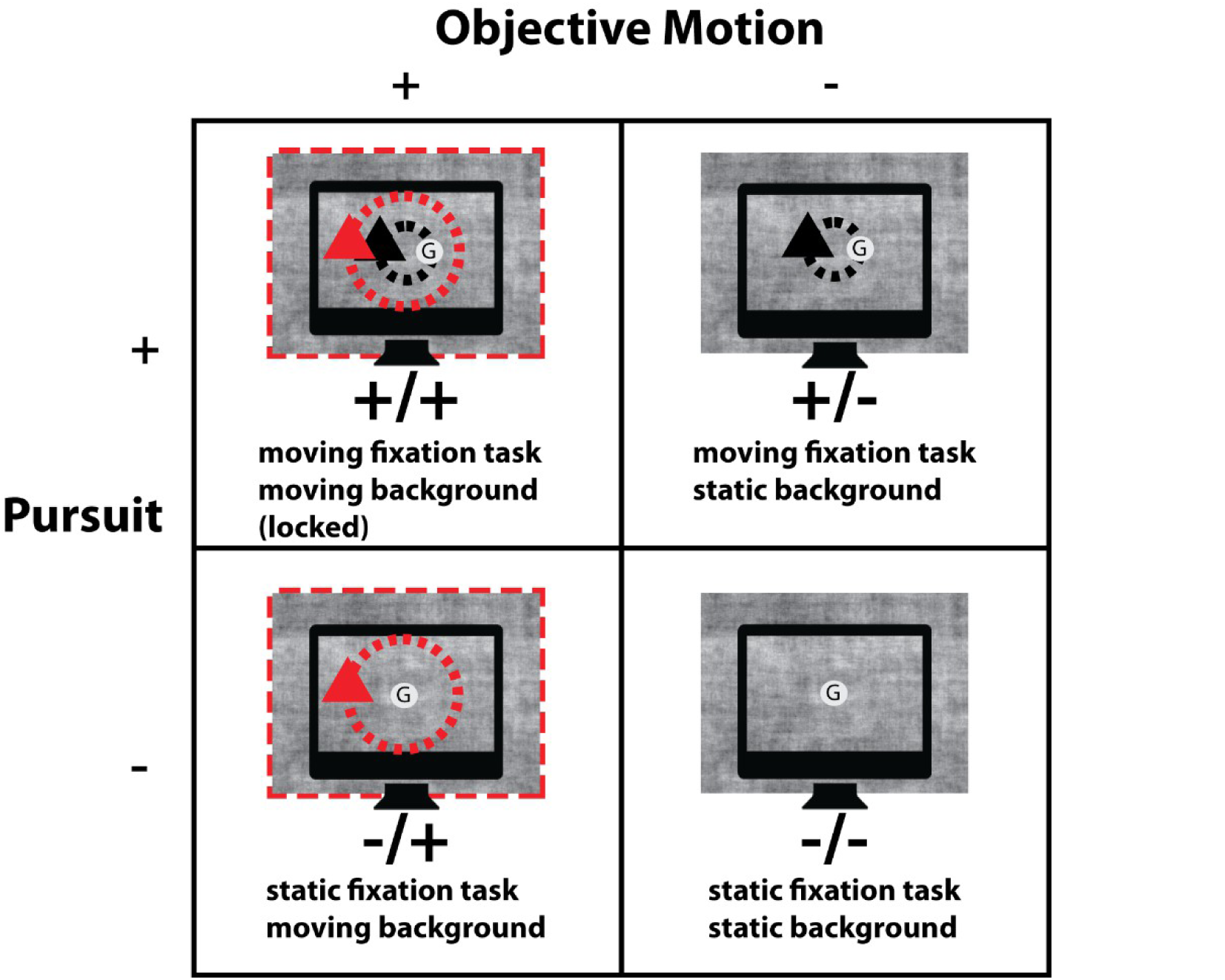
Four stimulus conditions presented in the main experiment, which were produced from a 2 x 2 factorial design with factors “pursuit” (on/off) and “objective motion” (on/off). The grey fixation disk with a one-back character-matching task was present at all times. Motion was planar on a circular trajectory. Movement of the fixation task resulted in pursuit whereas movement of the background on the same trajectory produced objective motion. The four conditions were as follows: +/+: moving fixation task with moving background (both trajectories were locked), −/+: static fixation task with moving background, +/−: moving fixation task with static background and −/−: both fixation task and background are static.

### FMRI Data Preprocessing

Data were analysed using SPM8 (www.fil.ion.ucl.ac.uk/spm/) and MATLAB 8.4 (2014b). Preprocessing steps were as follows: the functional images were realigned and the anatomical image was coregistered to mean functional image. The anatomical image was also normalized to MNI space. Functional images were smoothed with a 4 mm full-width at half maximum Gaussian kernel.

### Statistical Analysis

After preprocessing, each run of each participant (corresponding to one speed level) was analysed separately using the general linear model (GLM) approach. Regressors included one regressor for each of the four conditions, and one regressor for button presses. In addition, the following regressors of no interest were included: six motion realignment regressors, and an additional regressor of global mean signal, which was orthogonalized with respect to the conditions of interest in the design matrix (<u>Desjardins, Kiehl, & Liddle, 2001; Van Dijk et al., 2010</u>). High pass filtering with 128 s cut-off value was applied.

For each participant and each ROI, beta values were extracted for each condition and each speed level. For each participant, the resulting 24 (4 x 6) beta-values per ROI were z-normalized prior to contrast calculations and random effects ANOVA calculations: the mean was subtracted, followed by division by the standard deviation. This yielded 24 z-values for each participant and each ROI.

For each of the repeated measures ANOVAs conducted, Mauchly’s sphericity test was applied and in case of violation of sphericity, Greenhouse-Geisser correction was used.

### Statistical Contrasts

‘Objective motion’ was defined as the contrast of both conditions containing background motion versus both that did not, i.e. ((−/+) plus (+/+)) versus ((+/−) plus (−/−)). Note that pursuit was matched (equally present on either side).

‘Retinal motion’ was defined as both conditions containing retinal motion versus both that did not ((-/+) plus (+/-)) versus ((+/+) plus (-/-)). Again pursuit was matched.

‘Motion diff’: the *difference* between objective and retinal motion boils down to ((+/+) versus (+/−)) that again has matched pursuit conditions. This difference reliably and near-exclusively activates V3A in every participant (<u>Fischer et al., 2012a</u>).

‘Motion sum’ is the sum of objective and retinal motion ((−/+) versus (−/−)). This equals moving versus static background during fixation, and corresponds to the contrast used by previous motion studies, including those on speed processing. The responses to this contrast were calculated only for visualization and as a reference.

‘Pursuit’ was defined as ((+/+) plus (+/−)) versus ((−/+) plus (−/−)). This contrast contains several poorly controlled contributing factors, i.e. in addition to pursuit-controlling neural mechanisms also peripheral retinal motion beyond the projection screen that is not controlled, and a higher fixational error compared to non-pursuit (<u>Fischer et al., 2012a</u>). This contrast was therefore not analysed further, but was included in illustrations as a reference.

The above contrasts were calculated for each speed separately, using the z-normalized beta-values.

### Eye Tracking

An infrared camera based eye tracker (Eye-Trac 6; Applied Science Laboratories) was used in order to record eye position during the experiment together with Viewpoint Eyetracker software (Arrington Research, Scottsdale, USA) with a sampling rate of 60 Hz. Eye tracking data was analysed by the following steps: preprocessing was done by removing blinks and smoothing with a 200 ms running average window. The data was grouped according to the conditions. Root mean square error (RMSE) of eye position relative to the fixation disk was used to calculate fixation accuracy. This calculation was done separately for each condition, participant and speed. The same tests as conducted for the ROI data were applied to eye-tracking data.

## Results

We explored the responses to 6 different levels of speed, separately for objective motion and retinal motion, in independently localized motion responsive ROIs V3A, V6, V5/MT, MST and CSv.

For reference, Figure 2 plots values for the sum of objective and retinal motion. The sum (i.e. moving versus static background during fixation) corresponds to the contrast used in previous studies on motion and speed processing (e.g. (<u>Chawla et al., 1999; Chawla et al., 1998</u>).

**Figure 2.**
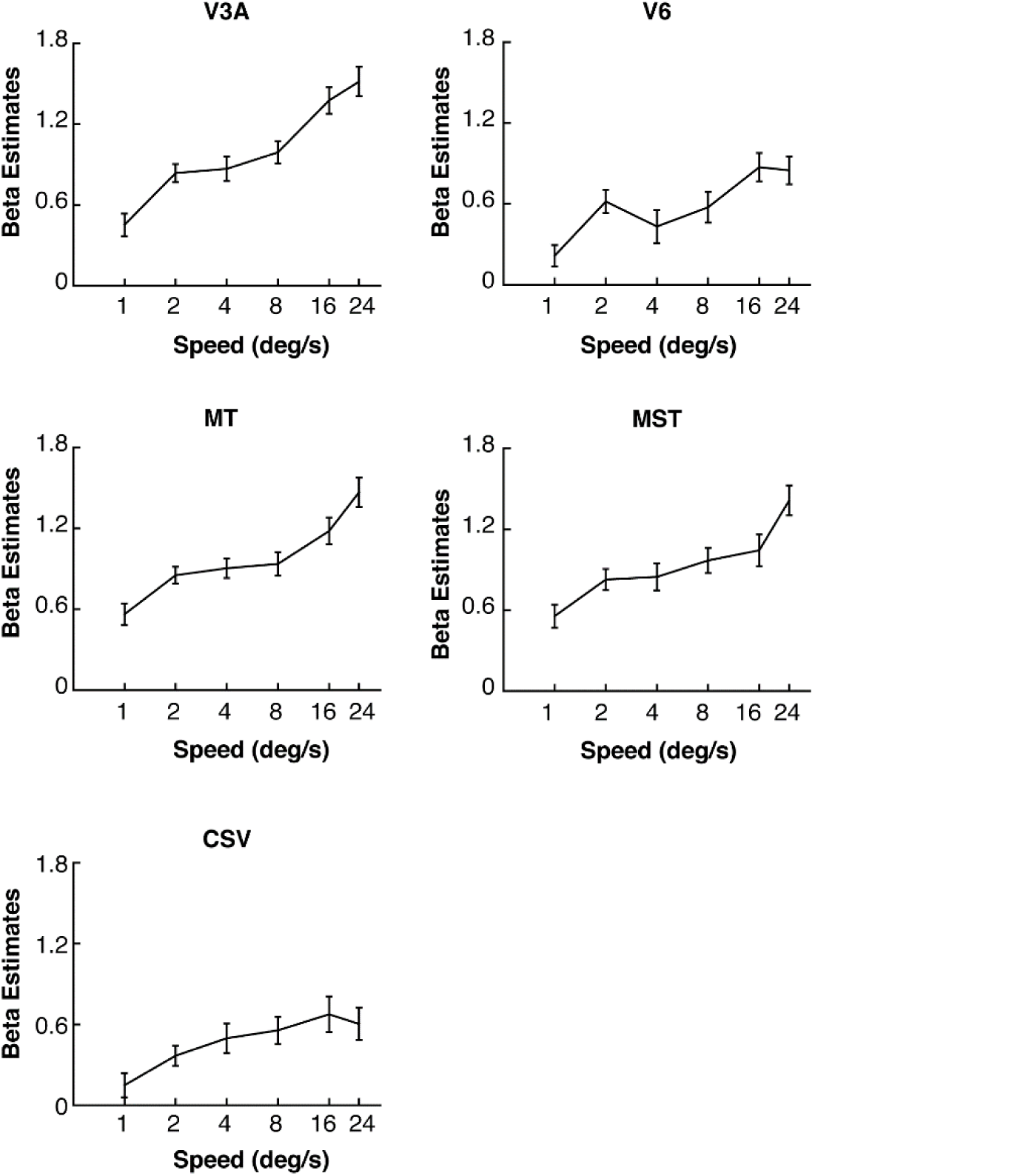
Contrast for ‘motion sum’. (−/+) vs. (−/−), as a function of speed. Here, values for moving background vs. static during fixation as a function of speed are shown for each ROI separately. This contrast would result in the sum of objective and retinal motion and was used in previous studies. Note that all data are z-normalized. Error bars show standard error of mean (SEM).

Figure 3 plots contrast values for objective and retinal motion (see methods for contrast definitions) as a function of speed. Contrast values were calculated for each speed separately using z-normalized beta-values (see methods). The difference between objective and retinal motion, i.e. preference for objective over retinal motion, has previously been shown to strongly activate V3A (<u>Fischer et al., 2012a</u>).

**Figure 3.**
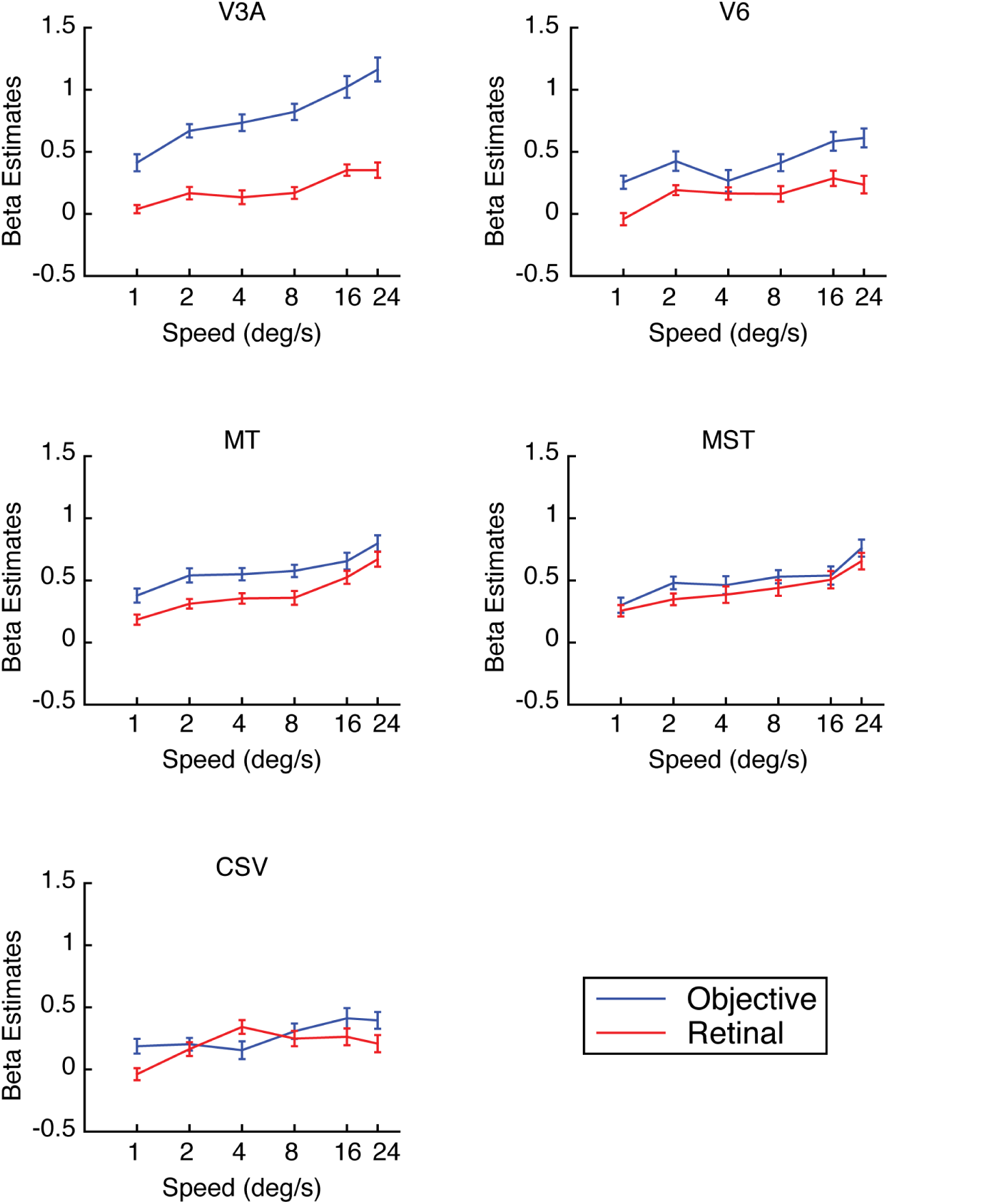
Contrast values for objective and retinal motion as a function of speed. Note that all data are z-normalized. Error bars show standard error of mean (SEM).

The results plotted in Figure 3 suggest three main findings. First, all regions tended to increase their responses with increasing speeds for both, objective and retinal motion. Second, V3A, V6 and to a lesser extent also V5/MT and MST had a higher overall response (i.e. offset) of objective relative to retinal motion. Third, the *slope* of the objective and retinal speed responses differed in V3A, with a steeper slope for objective motion.

In order to quantify these raw observations, we modelled each response using a simple linear fit to obtain a measure for the slope (b) and one for the offset (a), i.e. y = b*x +a, with y being the normalized contrast values, and x being the speed and it was scaled between −0.5 and +0.5. Previous single-unit recording studies for V5/MT reported a logarithmic speed tuning in motion responsive regions (<u>Nover et al., 2005</u>; <u>Priebe et al., 2003</u>), and, compatible with those, prior fMRI speed tuning studies also found log-like tuning in the speed range tested here (<u>Chawla et al., 1999</u>).

We hence directly tested two regression models: one modelled linear speed levels (x-axis: 1, 2, 4, 8, 16, 24 deg/s) and the other logarithmic scaling of speed (x-axis: 0, 1, 2, 3, 4, 4.585). We then tested which one provided a better fit by calculating the correlation coefficient for each, and comparing them in paired t-tests (after Fischer-Ztransforming r-values to z-values) for each ROI across participants (Table 1).

**Table 1:** Results of t tests comparing the fitness of regression models between linear scale and logarithmic scale of speed. For objective motion, retinal motion and pursuit, separately, we fitted regression models using either linear scale (1, 2, 4, 8, 16, 24 deg/s) or logarithmic scale (0, 1, 2, 3, 4, 4.585 deg/s) for speed. Next, for each ROI and motion type, we compared the correlation coefficients using t tests. P values that are significant after Bonferroni correction are shown in bold.

| p values | CSV | V3A | V5/MT | MST | V6 |
| --- | --- | --- | --- | --- | --- |
| objective | 0.77 | 0.328 | 0.157 | 0.155 | 0.189 |
|  | (t(37) = -0.3) | (t(37) = -1) | (t(39) = -1.4) | (t(37) = -1.5) | (t(37) = -1.3) |
| retinal | 0.322 | 0.823 | 0.964 | 0.984 | **0.00056** |
|  | (t(37) = 1.) | (t(37) = -0.23) | (t(39) = -0.05) | (t(37) = -0.02) | (t(37) = 3.8) |
| pursuit | 0.77 | 0.027 | 0.326 | 0.525 | 0.954 |
|  | (t(37) = 0.3) | (t(37) = 2.3) | (t(39) = 1) | (t(37) = 0.6) | (t(37) = 0.06) |

After applying Bonferroni correction for 15 comparisons, the only significant result was for V6, showing that retinal motion was fitted better by logarithmic scaling. Judging from the low t-values for all other tests, there was no clear preference for either type of scaling. We used logarithmic scaling of speed for the rest of the analysis.

Next, we tested regression slopes for each motion type and every ROI against zero, and compared regression slopes between objective and retinal motion for each ROI using paired t-tests. Independently of this, we also tested the mean signal (offset) in the same way. Figure 4A shows the regression slopes for objective and retinal motion for all ROIs, figure 4B mean signals. All t-test results were corrected for 15 comparisons using Bonferroni-Holm correction. As shown in table 2, the slope for objective motion was significantly different than zero for all, whereas for retinal motion, it was only significant for V5/MT, MST and V3A but not for Csv or V6. The comparison of objective and retinal motion regression slopes was only significant for V3A. As shown in table 3, mean responses to objective and retinal motion were significant for all ROIs and V3A, V5/MT and V6 had significantly higher mean responses to objective motion than retinal motion.

**Table 2.** Paired t-test results for slopes

| p values | CSV | V3A | V5/MT | MST | V6 |
| --- | --- | --- | --- | --- | --- |
| **Objective slope** | **0.0001**<br>(t(37)=4.33) | **$0.3 * 10^{-9}$**<br>(t(37)=8.38) | **$0.2 * 10^{-6}$**<br>(t(39)=6.26) | **$0.2 * 10^{-5}$**<br>(t(37)=5.59) | **$0.2 * 10^{-4}$**<br>(t(37)=4.87) |
| **Retinal slope** | 0.005<br>(t(37)=2.95) | **$0.2 * 10^{-4}$**<br>(t(37)=4.87) | **$0.7 * 10^{-7}$**<br>(t(39)=6.61) | **$0.2 * 10^{-4}$**<br>(t(37)=4.93) | 0.004<br>(t(37)=3.03) |
| **Objective slope**<br>**vs.**<br>**retinal slope** | 0.59<br>(t(37)=0.54) | **0.0009**<br>(t(37)=3.58) | 0.24<br>(t(39)=-1.19) | 0.99<br>(t(37)=-0.001) | 0.4<br>(t(37)=0.85) |

**Figure 4:**
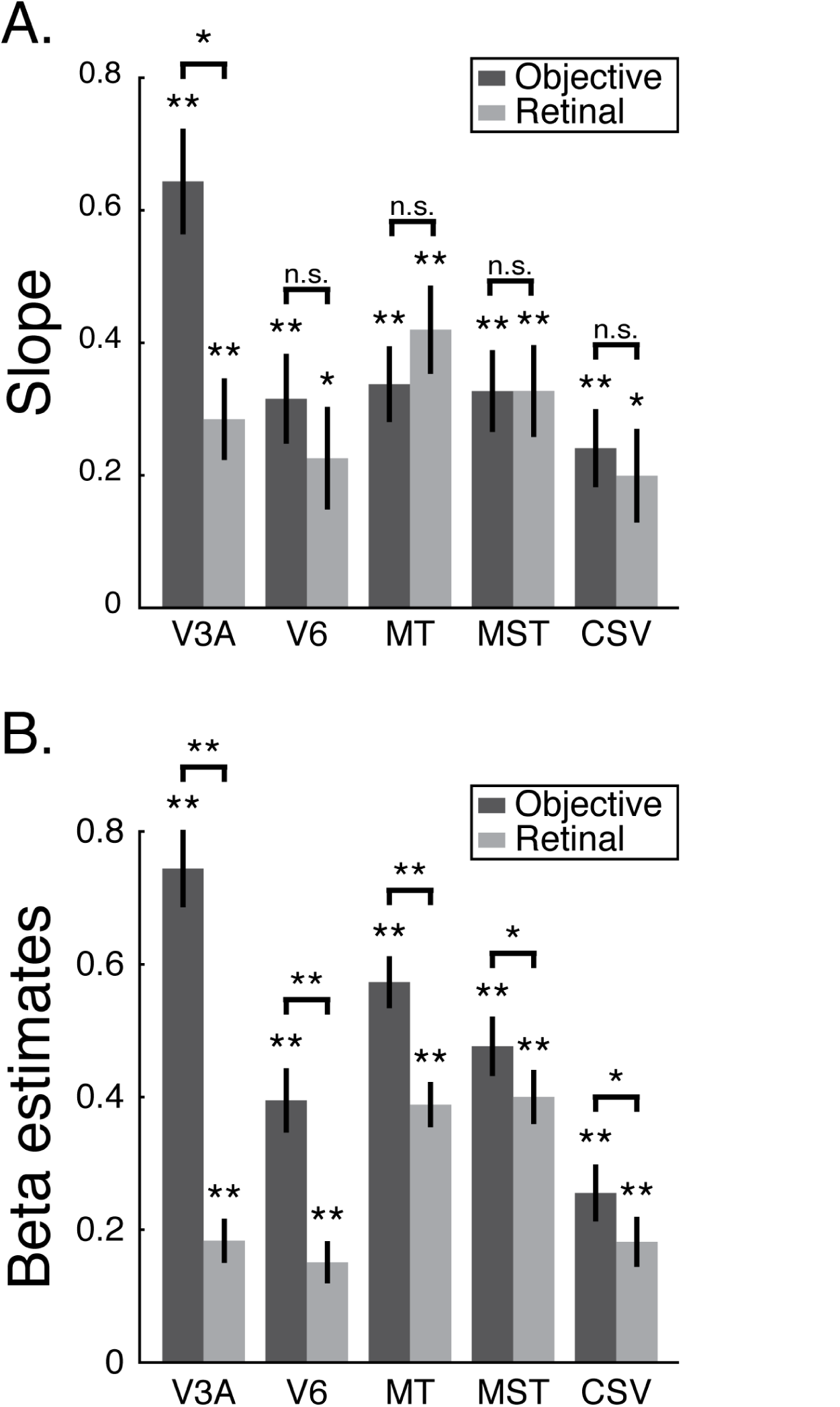
Regression slopes and mean signal for objective and retinal motion. (A) Regression slopes calculated for objective and retinal motion, for each ROI separately. Every ROI was modulated by both objective motion and retinal motion speed. Only in V3A, objective motion speed slope was significantly higher than the slope of retinal motion speed. (B) Mean signal (offset) for objective and retinal motion. For every ROI, the mean objective motion and mean retinal motion was significantly greater than zero and mean objective motion was significantly higher than mean retinal motion. Note that all data are z-normalized. ** p<0.001, * p<0.05, Bonferroni-Holm corrected. Error bars show standard error of mean (SEM).

**Table 3.**
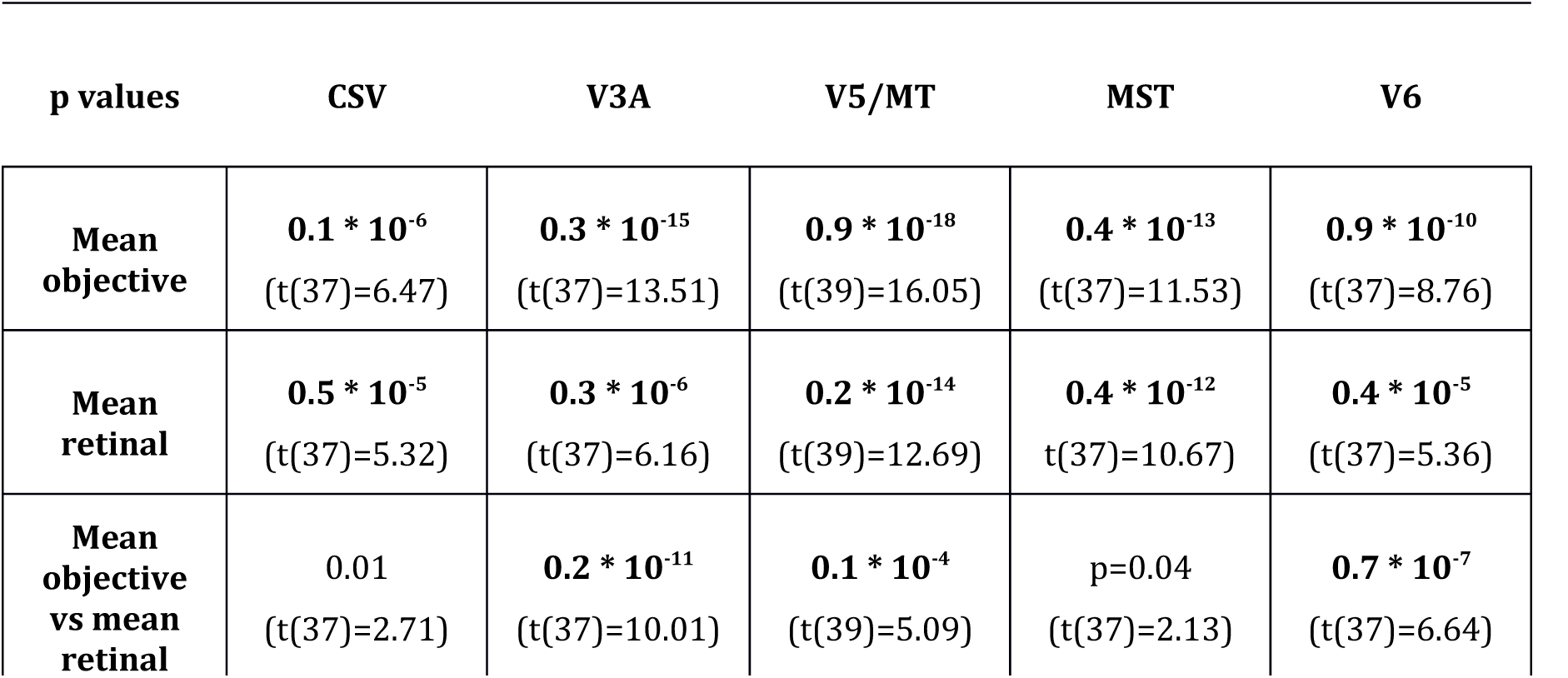
Paired t-test results for mean responses

### Behavioral Data

During the experiment, participants were required to perform a character back-matching task at all times. Average rate of correct responses was 0.83 ± 0.04 (mean ± std) and mean response time was 0.52 ± 0.12 s (mean ± std). Similar to fMRI data analysis, for each participant, we first calculated correct response rate during objective motion and retinal motion separately. Next, for each participant, we fit two separate GLMs to the correct response data with 2 regressors each; one for the speed and the other being all ones. The speed regressor was first calculated in logarithmic scale (0, 1, 2, 3, 4, 4.585 deg/s) and then scaled between −0.5 and +0.5. This way the first regressor modelled the slope while the second one was modelling for the mean correct response. For each participant, we calculated slope of correct responses during objective and retinal motion speeds separately.

Using paired t-tests we tested the correct response rate slopes during objective and retinal motion. We also did additional t-tests to compare objective and retinal motion slopes. None of the test results reached significance (objective motion: t (19) = 1.41, p = 0.176; retinal: t (19) = 0.08, p = 0.936; objective vs. retinal: t (19) = 0.54, p = 0.594). The slopes for task responses during objective and retinal motion speed are shown in figure 5A.

**Figure 5.**
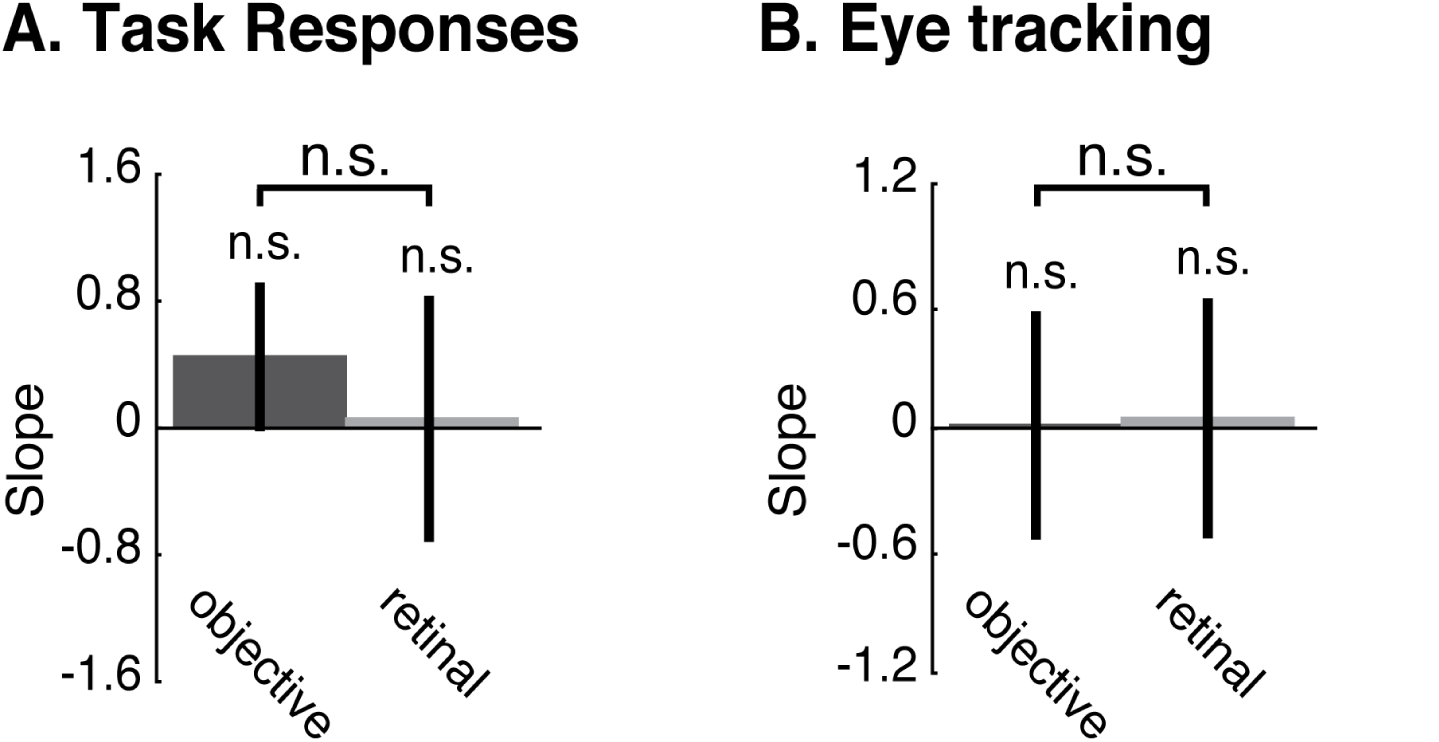
Regression slopes for behavioral and eye tracking data. (A) Regression slopes calculated for correct task responses during objective motion speed contrast and retinal motion speed contrast separately. There is no significant effect of speed during retinal and objective motion and there is no significant difference between task performance during objective and retinal motion speed. (B) Regression slopes calculated for RMSE of eye position relative to the fixation disk during objective motion speed and retinal motion contrast separately. Here, there is no significant effect of objective motion speed or retinal motion speed on RMSE of eye position and there is no significant difference between objective and retinal motion speed on RMSE of eye position. Error bars show standard error of mean (SEM).

### Eye Tracking Data

We collected eye-tracking data for 13 of the 20 participants. After preprocessing of the data, we calculated RMSE of eye position relative to the fixation disk and used this for comparing fixation accuracy across conditions and different levels of speed. The mean RMSE ± SEM across participants for each speed and condition is shown in table 4.

**Table 4.** RMSE of eye position for each condition and speed‥ Values illustrate mean ± standard error of mean (SEM)

| Conditions | 1 deg/s | 2 deg/s | 4 deg/s | 8 deg/s | 16 deg/s | 24 deg/s |
| --- | --- | --- | --- | --- | --- | --- |
| (+/+) | 2.24 ± 0.30 | 2.02±0.21 | 2.13±0.21 | 2.44±0.29 | 2.83±0.19 | 3.65±0.15 |
| (-/ +) | 1.35 ± 0.21 | 1.57±0.18 | 1.57±0.24 | 1.28±0.22 | 1.43±0.25 | 1.49±0.19 |
| (+/-) | 2.01 ± 0.22 | 2.26±0.25 | 2.28±0.27 | 2.25±0.21 | 2.86±0.20 | 3.56±0.10 |
| (-/-) | 1.75 ± 0.31 | 1.31±0.18 | 1.66±0.25 | 1.23±0.18 | 1.57±0.29 | 1.44±0.22 |

For each participant, using RMSE of eye position, we calculated regression slopes for objective and retinal motion speed separately. We calculated RMSE of eye position during objective and retinal motion for each participant separately and fit GLMs to calculate slopes, similar to fMRI and behavioral data analyses.

Next, we did paired t tests to test the significance of objective and retinal motion slopes as well as to compare objective and retinal motion slopes. None of the t-tests yielded a significant difference (objective motion: t (12) = 1.13, p = 0.28; retinal motion: t (12) = 0.18, p = 0.859; objective vs. retinal motion: t (12) = 0.56, p = 0.589). The slopes for eye position during objective and retinal motion are shown in figure 5B.

## Discussion

Many motion processing regions contain neurons that differentiate between world-centered (objective) and eye-centered (retinal) motion (<u>Dicke, Chakraborty, & Thier, 2008</u>; <u>Erickson & Thier, 1991</u>; <u>Galletti et al., 1990</u>; <u>Ilg et al., 2004</u>; <u>Zhang, Heuer, & Britten, 2004</u>). Correspondingly, recent fMRI studies showed that net BOLD responses in human motion regions can vary strongly by their preferences to both types of motion (<u>Arnoldussen et al., 2011</u>; <u>Fischer et al., 2012a</u>). Speed dependency of either type of motion has never been examined before. Prior speed studies (that measured the sum of both motion types) are over 15 years old, used only 3 participants, and no distraction task, and may contain attention-speed interactions, and potentially low-level motion-streak effects due to older projection technology (<u>Chawla et al., 1999; Chawla et al., 1998</u>). Finally, speed tuning is entirely unknown for several regions that only recently came into the spotlight of human fMRI research, such as CSv and V6.

Hence, in this study, we re-addressed the question of speed dependent responses across independently localized visual motion areas V3A, V6, V5/MT, MST and CSv, in 20 participants. We used stimuli with natural image statistics in order to stimulate the motion system with stimulus properties it evolved for. In addition, we used a pursuit paradigm to address this question that enabled us to separate objective and retinal motion while matching eye movements across both types of motion (<u>Fischer et al., 2012a</u>). Finally, we used a projector with 120 Hz refresh rate that had been specifically selected for low motion-streak effects.

We found that all regions increased responses monotonically as a function of speeds for both, retinal and objective motion, for speeds between 1-24 deg/s. In contrast to prior fMRI studies, we did not observe an inverted U shape response for either type of motion. This was most likely due to our upper limit of 24 deg/s, whereas electrophysiology studies show that in many motion regions optimal speed tuning is between 32 deg/s or far beyond. We were constrained here to 24 deg/s, as beyond this, reliable pursuit performance would have been difficult to achieve.

The responses in all regions were fit well using either a linear or logarithmic speed-tuning function. The slopes of the speed-response functions did not differ between objective and retinal motion in all ROIs, except for V3A. V3A stood out in that its objective motion slope was significantly higher than that for retinal motion. In V3A, also the mean response to objective motion was by far higher than that to retinal motion, with a significant difference also being present in V6, V5/MT, and MST and CSv, in descending order.

One could argue that the speed responses here could have resulted from eye movement related biases between conditions. It is hence important to note that for both, objective and retinal motion, pursuit-versus-non-pursuit conditions were entirely matched and cancelled out. Moreover, our eye tracking and behavioral data results show no confound; there were no effects of speed during objective or retinal motion.

To our knowledge this is the first systematic study of speed tuning conducted using modern imaging standards, eye-tracking, a demanding attention task, and including human motion regions V6 and CSv. In addition, it is the first to segregate objective and retinal motion speed tuning.

### Speed Encoding in V5/MT and MST

In this study, V5/MT and MST showed significant speed tuning to both objective and retinal motion. Moreover, neither V5/MT nor MST seemed to differentiate between speed of objective motion and speed of retinal motion in terms of response slopes, but both regions had marginally higher overall responses to objective motion.

As shown by single neuron recordings and lesion studies, motion responsive regions V5/MT has a role in speed perception and processing of speed related motion signals as most MT neurons are speed responsive (<u>Dursteler & Wurtz, 1988</u>; <u>Krekelberg et al., 2006</u>; <u>Liu & Newsome, 2005</u>; <u>Maunsell & Van Essen, 1983b</u>; <u>Newsome et al., 1985</u>; <u>Orban, Saunders, et al., 1995</u>; <u>Pasternak & Merigan, 1994</u>; <u>Perrone & Thiele, 2001</u>; <u>Yamasaki & Wurtz, 1991</u>). Moreover, different studies reported different preferred range of speed in monkey V5/MT neurons, for instance between 2 and 256 deg/s (<u>Maunsell & Van Essen, 1983b</u>) or 5-250 deg/s (<u>Rodman & Albright, 1987</u>) while the optimal speed is shown to be around 32 deg/s in many studies (<u>Cheng, Hasegawa, Saleem, & Tanaka, 1994</u>; <u>Mikami, Newsome, & Wurtz, 1986</u>; <u>Newsome, Mikami, & Wurtz, 1986</u>). Our results therefore fit to the previous findings since we do not see any optimal range of speed and only observe monotonic increase in speed responses when tested between 1 to 24 deg/s. Testing higher speed ranges might provide the optimal speed range in human.

Similarly, single neuron studies in monkey MST also showed speed tuning in these neurons (<u>Duffy & Wurtz, 1997</u>). These authors tested speed tuning of MST neurons for radial and circular motion types and found that in about 2/3rd of the tested neurons, responses were modulated by speed when tested for speed range between 10-80 deg/s (<u>Duffy & Wurtz, 1997</u>). Previous studies reported that MST neurons preferred high stimulus speeds both during laminar motion and optic flow (<u>A. K. Churchland & Lisberger, 2005</u>; <u>Duffy & Wurtz, 1997</u>; <u>Kawano, Shidara, Watanabe, & Yamane, 1994</u>; <u>Orban, Lagae, Raiguel, Xiao, & Maes, 1995</u>; <u>Tanaka & Saito, 1989</u>). While Orban and colleagues reported the optimal speed range of MST neurons between 15-20 deg/s (<u>Orban, Lagae, et al., 1995</u>), Duffy and colleagues found that approximately 40%of the neurons tested showed increasing response profile with increasing speed (<u>Duffy & Wurtz, 1997</u>). Additionally, Kawano and colleagues reported the optimal speed for MST single neuron responses as 160 deg/s (<u>Kawano et al., 1994</u>). These results reveal that while there seems to be no consensus on optimal speed range of MST neurons, it is possible that similar to the case in V5/MT, we also did not reach optimal speed range in MST.

Single neuron studies showed that MT processes retinal signals whereas MST responses take both retinal and extraretinal signals into account (<u>Erickson & Thier, 1991</u>; <u>Galletti & Fattori, 2003</u>; <u>Ilg & Thier, 2003</u>). In monkey, a large fraction of neurons in MSTd have been shown to compensate for speed of pursuit during eye movements whereas neurons in V5/MT are generally more responsive to retinal motion (<u>Chukoskie & Movshon, 2009; Inaba, Miura, & Kawano, 2011</u>; <u>Lee, Pesaran, & Andersen, 2007</u>; <u>Shenoy, Crowell, & Andersen, 2002</u>). For instance, MSTd neurons were shown to compensate for pursuit speed when tested for pursuit with the speed of 2.58, 5.05 and 9.22 deg/s (<u>Shenoy et al., 2002</u>). Moreover, the same study showed that MSTd uses both retinal and extraretinal input to compensate for pursuit, since the preferred pursuit and retinal motion directions are opposite of each other (<u>Shenoy et al., 2002</u>). Similarly, Lee and colleagues found that MSTd neurons also compensate for changes in translation speed (<u>Lee et al., 2007</u>).

### Perceptual effects

The question thus arises why here V5/MT showed higher mean objective motion compared to retinal motion. One possible account may have to do with perceptual effects, which, in contrast to spiking activity, are often dominantly reflected in fMRI signal (<u>Bartels, Logothetis, & Moutoussis, 2008</u>). In humans, V5/MT+, as well as V3A, was previously shown to have a role in speed perception (<u>McKeefry et al., 2008</u>) and V5/MT+ was shown to have headcentric speed tuning during optic flow stimuli (<u>Arnoldussen et al., 2011</u>). Also, previous electrophysiology studies showed that MT responses correlate with speed perception (<u>Krekelberg & van Wezel, 2013</u>; <u>Liu & Newsome, 2005</u>). Additionally, the speed of background motion during eye fixation has been shown to be perceived higher than the speed of retinal motion of smooth pursuit eye movements on a static background, even though the retinal speed induced in both conditions are exactly the same (Aubert-Fleischl effect) (<u>Aubert, 1886</u>; <u>Dichgans, Körner, & Voigt, 1969</u>). Regarding this, it is possible that in the present study, the speed during objective and retinal motion conditions were perceived differently, meaning the speed during (-/+) condition was perceived higher than the speed during (+/-) condition. Additionally, the visual system is thought to be more tolerable to retinal motion in the opposite direction of the pursuit, since this type of motion is natural during pursuit and could be arising from the eye movements itself and less tolerable to retinal motion in the same direction as pursuit, as this could indicate ‘real’ object motion (<u>Lindner, Schwarz, & Ilg, 2001</u>). This was shown by previous studies; when brief injections of background motion was introduced during smooth pursuit eye movements, it only caused an increase in eye velocity when it is in the same direction as pursuit, but did not cause any increase when it is in opposite direction (<u>Lindner et al., 2001</u>). Additionally, V5/MT and MST are densely connected to each other and MST is higher in the visual processing hierarchy than V5/MT (<u>Boussaoud, Ungerleider, & Desimone, 1990</u>; <u>Maunsell & Van Essen, 1983a</u>; <u>Ungerleider & Desimone, 1986</u>). So, another possibility is that feedback from MST to V5/MT could be resulting in higher objective motion responses compared to retinal motion responses in V5/MT.

### Relation to prior fMRI studies

Two fMRI studies showed that when combined objective/retinal motion speed was used with the range of speed between 3.7 and 61.6 degrees and between 1 and 32 degrees, V5/MT (and V3A) showed non-linear (inverted U shape) speed responses and their optimal speed range was between 7 deg/s and 30 deg/s and between 4 deg/s and 8 deg/s in two separate studies (<u>Chawla et al., 1999; Chawla et al., 1998</u>).

However, our results disagree with the optimal speed ranges reported by those studies from Chawla and colleagues (<u>Chawla et al., 1999; Chawla et al., 1998</u>). According to their reports, speed range used in the present study, which is between 1-24 deg/s, should be more or less the optimal range to result in inverted U-shape speed responses, similar to their findings. Although some previous single neuron studies report a similar inverted U-shaped speed response, the range of speeds reported in those studies are much wider and for the majority of cells, optimal speed is considerably higher than those reported by Chawla and colleagues (<u>Chawla et al., 1999; Chawla et al., 1998</u>; <u>Cheng et al., 1994; Mikami et al., 1986</u>; <u>Rodman & Albright, 1987</u>). One can think the stimuli used as one main difference. In both mentioned studies, Chawla and colleagues used dots moving radially from centre towards the edges of the screen (optic expansion flow) (<u>Chawla et al., 1999; Chawla et al., 1998</u>). However, they do not report how many dots were implemented or whether new dots appeared at the centre of the screen. Inherent to any expansion flow display is that dot-appearances, typically foremost near the centre of the screen, scale with speed. In this case, the inverted-U shape could simply be an artefact of temporal frequency tuning. It is well known that V5/MT is responsive to flickering static stimuli (<u>Malonek, Tootell, & Grinvald, 1994; Tootell et al., 1995</u>). Further, in a previous fMRI study, Singh and colleagues (<u>Singh, Smith, & Greenlee, 2000</u>) investigated spatial and temporal frequency tuning of visual areas in human brain and found a very similar inverted-U shape response tuning for temporal frequency. Additionally, those studies conducted by Chawla et al. did not have any attention task (apart from fixation) and this could lead to potential attentional load differences across speed (<u>Chawla et al., 1999; Chawla et al., 1998</u>). Previously, attention to speed was shown to strengthen the responses in V5/MT (<u>Beauchamp et al., 1997</u>). One should also keep in mind that mere data quality, field-strength, and standards for number of participants (3 participants vs. 20) have changed dramatically (<u>Chawla et al., 1999; Chawla et al., 1998</u>).

### Speed Encoding in V3A

It is well known that V3A is a part of motion processing network (<u>Galletti et al., 1990; Tootell et al., 1997</u>). In humans, V3A has the second highest motion responses after V5/MT and motion responses in human V3A are more similar to that of monkey V3, not V3A (<u>Tootell et al., 1997</u>). However, speed tuning of V3A is not extensively studied for different types of motion. In humans, attention to speed of motion activated V3A (<u>Sunaert, Van Hecke, Marchal, & Orban, 2000</u>). In the present study, V3A responses show speed tuning for both objective and retinal motion types. These results are in accord with previous studies that showed that V3A have a crucial role for perceiving stimulus speed (<u>McKeefry et al., 2008; Pitzalis, Strappini, De Gasperis, Bultrini, & Di Russo, 2012</u>). It is possible that we explored only the lower end of the speed range, since V3A neurons in macaque monkey are shown to be sensitive to a wide range of speeds and they are even activated at speeds higher than 50 deg/s (<u>Galletti et al., 1990</u>).

Previous studies on single neuron responses showed that V3A neurons showed real-motion preference (<u>Galletti et al., 1990</u>) and approximately half of the V3A neurons in monkey were found to be gaze dependent (<u>Galletti & Battaglini, 1989</u>). In this study, V3A was the only region to significantly differentiate between the speed of objective and retinal motion. Moreover, V3A prefers objective motion to retinal motion at all speeds. This is compatible with previous studies about real motion responses in V3A (<u>Fischer et al., 2012a; Galletti & Fattori, 2003</u>). Even during 3D motion, V3A has been shown to have strong self-motion responses and encodes headcentric speed of rotation (<u>Arnoldussen et al., 2011</u>, <u>2015</u>). We conclude that during 2D motion, V3A is speed tuned to both objective and retinal motion, and it can differentiate between the speed of objective motion and retinal motion.

### Speed Responses in V6

V6 is a motion responsive region with large-field responses, and in particular responds to egomotion compatible motion such as 3D flow (<u>Cardin & Smith, 2011; Fattori, Pitzalis, & Galletti, 2009; Pitzalis et al., 2006; Pitzalis et al., 2013; Pitzalis et al., 2010</u>). V6 neurons in macaque respond to a wide variety of speeds, reported between 0 deg/s to 900 deg/s (<u>Galletti, Fattori, Battaglini, Shipp, & Zeki, 1996</u>) and a previous study reported that it showed higher responses for the fast speed when tested for 3 deg/s and 25 deg/s (<u>Pitzalis et al., 2012</u>). Further, Arnoldussen et al. (<u>Arnoldussen et al., 2011</u>) showed that V6, together with V3A and MT+, is encoding headcentric speed of motion. We found that V6 showed speed-tuned responses to objective motion, and to a lesser extent to retinal motion and that it did not show differential responses between objective and retinal motion speed tuning.

Similar to V3A, V6 is also responsive to real motion (<u>Fischer et al., 2012a; Galletti & Fattori, 2003</u>). V6 is thought to be encoding extrapersonal space since majority of neurons in macaque V6 are reported to be eye position sensitive and some neurons are reported to be ‘real-position’ cells (<u>Galletti, Battaglini, & Fattori, 1995</u>). In the present study, we used 2D planar motion, whereas most of the previous studies mentioned here used 3D coherent motion. When 2D planar motion was used, V6 responses were suppressed during retinal motion (<u>Fischer et al., 2012a</u>). Objective motion preference found in V6 in this study is in accord with egocentric / headcentric motion preference of V6.

In humans, V6 is shown to prefer near field stimuli (<u>Quinlan & Culham, 2007</u>). Together with this and its connections to V6A and other parietal regions that have a role in grasping and to visual cortex (<u>Galletti et al., 2001</u>), V6 is thought to have a role in encoding the movement of graspable objects (<u>Galletti & Fattori, 2003; Galletti et al., 2001</u>). Being able to differentiate objective and retinal motion, as well as preference for high speeds, can both be explained by the preference for near visual field and encoding of graspable object motion. Objects that are closer to us seem to move faster than objects that are further away. Thus, the higher speed preference in V6 is in line with its role in encoding for graspable objects that are in peripersonal space.

### Speed Responses in CSv

CSv is a recently defined visual region in dorsal posterior cingulate sulcus (dPCC) that is responsive to complex motion and it is specialized in processing self motion related signals and parsing optic flow (<u>Antal, Baudewig, Paulus, & Dechent, 2008; Cardin & Smith, 2010; Fischer et al., 2012b; Wall & Smith, 2008</u>). Although there are studies in macaque posterior cingulate sulcus showing visual responses (<u>Dean, Crowley, & Platt, 2004</u>), it is not clear whether there is a homologue of human CSv in primates. Thus, there are not so many electrophysiology studies that are directly comparable.

Previously, CSv has been shown to monitor eye position during saccades and smooth pursuit eye movements (<u>Olson, Musil, & Goldberg, 1996</u>). CSv was also shown to compensate for self-induced retinal eye movements (<u>Fischer et al., 2012b</u>). Our results regarding higher mean objective motion responses compared to mean retinal motion responses in CSv are consistent with these studies. A recent fMRI study, which was conducted using 3D optic flow motion, reported no significant speed tuning for retinal or headcentric motion in CSv, whereas there was a significant speed tuning response for pursuit (<u>Arnoldussen et al., 2011</u>). While our results seem to disagree with those findings, it is possible that eye position related signals used for compensation of eye movements could be involved differently in calculations regarding planar motion and more complex motion such as heading related optic flow signals (for a detailed discussion, please see (<u>Fischer et al., 2012a</u>)).

To our knowledge, this is the first study to systematically investigate the speed tuning in CSv. We found that CSv showed significant speed tuning for both objective motion and to a lesser extent for retinal motion, but its speed tuning did not differentiate between these two types of motion. Although the mean objective motion response in CSv was significantly higher than the mean retinal motion response, this was not consistent cross all speeds (Figure 3).

## Conclusion

In conclusion, our results provide the speed tuning of motion responsive regions during objective and retinal motion. Furthermore, V3A is the only motion responsive region that shows different speed tuning to objective and retinal motion. These results support the view that human V3A encodes primarily objective rather than retinal motion signals even during a range of motion speeds.

## Acknowledgements

This work was funded by the Centre for Integrative Neuroscience Tübingen through the German Excellence Initiative (EXC307) and by the Max Planck Society, Germany.

## References

Antal, A., Baudewig, J., Paulus, W., & Dechent, P. (2008). The posterior cingulate cortex and planum temporale/parietal operculum are activated by coherent visual motion. Visual neuroscience, 25, 17–26. doi: 10.1017/S0952523808080024

Arnoldussen, D. M., Goossens, J., & van den Berg, A. V. (2011). Adjacent visual representations of self-motion in different reference frames. Proceedings of the National Academy of Sciences of the United States of America, 108(28), 11668–11673. doi: 10.1073/pnas.1102984108

Arnoldussen, D. M., Goossens, J., & van den Berg, A. V. (2015). Dissociation of retinal and headcentric disparity signals in dorsal human cortex. Frontiers in Systems Neuroscience, 9, 16. doi: 10.3389/fnsys.2015.00016

Aubert, H. (1886). Die bewegungsempfindung. Pflügers Archiv European Journal of Physiology, 39(1), 347–370.

Bartels, A., Logothetis, N. K., & Moutoussis, K. (2008). fMRI and its interpretations: an illustration on directional selectivity in area V5/MT. Trends in neurosciences, 31, 444–453. doi: 10.1016/j.tins.2008.06.004

Bartels, A., Zeki, S., & Logothetis, N. K. (2008). Natural vision reveals regional specialization to local motion and to contrast-invariant, global flow in the human brain. Cerebral cortex, 18, 705–717. doi: http://dx.doi.org/10.1093/cercor/bhm107

Beauchamp, M. S., Cox, R. W., & DeYoe, E. A. (1997). Graded effects of spatial and featural attention on human area MT and associated motion processing areas. Journal of Neurophysiology, 78(1), 516–520.

Boussaoud, D., Ungerleider, L. G., & Desimone, R. (1990). Pathways for Motion Analysis-Cortical Connections of the Medial Superior Temporal and Fundus of the Superior Temporal Visual Areas in the Macaque. Journal of Comparative Neurology, 296(3), 462–495.

Brainard, D. H. (1997). The psychophysics toolbox. Spatial vision, 10(4), 433–436.

Brooks, J. L. (2012). Counterbalancing for serial order carryover effects in experimental condition orders. Psychological Methods, 17(4), 600–614. doi: 10.1037/a0029310

Cardin, V., & Smith, A. T. (2010). Sensitivity of human visual and vestibular cortical regions to egomotion-compatible visual stimulation. Cerebral cortex, 20(8), 1964–1973. doi: 10.1093/cercor/bhp268

Cardin, V., & Smith, A. T. (2011). Sensitivity of human visual cortical area V6 to stereoscopic depth gradients associated with self-motion. Journal of neurophysiology, 106(3), 1240–1249. doi: 10.1152/jn.01120.2010

Chawla, D., Buechel, C., Edwards, R., Howseman, A., Josephs, O., Ashburner, J., & Friston, K. J. (1999). Speed-dependent responses in V5: A replication study. Neuroimage, 9(5), 508–515.

Chawla, D., Phillips, J., Buechel, C., Edwards, R., & Friston, K. J. (1998). Speed-dependent motion-sensitive responses in V5: an fMRI study. Neuroimage, 7(2), 86–96.

Cheng, K., Hasegawa, T., Saleem, K. S., & Tanaka, K. (1994). Comparison of neuronal selectivity for stimulus speed, length, and contrast in the prestriate visual cortical areas V4 and MT of the macaque monkey. Journal of Neurophysiology, 71(6), 2269–2280.

Chukoskie, L., & Movshon, J. A. (2009). Modulation of visual signals in macaque MT and MST neurons during pursuit eye movement. Journal of neurophysiology, 102(6), 3225–3233. doi: 10.1152/jn.90692.2008

Churchland, A. K., & Lisberger, S. G. (2005). Discharge properties of MST neurons that project to the frontal pursuit area in macaque monkeys. J Neurophysiol, 94(2), 1084–1090.

Churchland, M. M., & Lisberger, S. G. (2001). Shifts in the Population Response in the Middle Temporal Visual Area Parallel Perceptual and Motor Illusions Produced by Apparent Motion. The Journal of neuroscience: the official journal of the Society for Neuroscience, 21(23), 9387–9402.

Dean, H. L., Crowley, J. C., & Platt, M. L. (2004). Visual and saccade-related activity in macaque posterior cingulate cortex. J Neurophysiol, 92(5), 3056–3068. doi: 10.1152/jn.00691.2003 00691.2003 [pii]

Desjardins, A. E., Kiehl, K. A., & Liddle, P. F. (2001). Removal of confounding effects of global signal in functional MRI analyses. Neuroimage, 13(4), 751–758. doi: http://dx.doi.org/10.1006/nimg.2000.0719

Dichgans, J., Körner, F., & Voigt, K. (1969). Vergleichende Skalierung des afferenten und efferenten Bewegungssehens beim Menschen: Lineare Funktionen mit verschiedener Anstiegssteilheit. Psychologische Forschung, 32(4), 277–295.

Dicke, P. W., Chakraborty, S., & Thier, P. (2008). Neuronal correlates of perceptual stability during eye movements. Eur J Neurosci, 27(4), 991–1002. doi: EJN6054 [pii] 10.1111/j.1460-9568.2008.06054.x

Duffy, C. J., & Wurtz, R. H. (1997). Medial superior temporal area neurons respond to speed patterns in optic flow. The Journal of neuroscience, 17(8), 2839–2851.

Dursteler, M. R., & Wurtz, R. H. (1988). Pursuit and optokinetic deficits following chemical lesions of cortical areas MT and MST. Journal of Neurophysiology, 60(3), 940–965.

Erickson, R. G., & Thier, P. (1991). A Neuronal Correlate of Spatial Stability during Periods of Self-Induced Visual-Motion. Experimental Brain Research, 86(3), 608–616.

Fattori, P., Pitzalis, S., & Galletti, C. (2009). The cortical visual area V6 in macaque and human brains. Journal of Physiology-Paris, 103(1–2), 88–97. doi: http://dx.doi.org/10.1016/j.jphysparis.2009.05.012

Fischer, E., Bulthoff, H. H., Logothetis, N. K., & Bartels, A. (2012a). Human areas V3A and V6 compensate for self-induced planar visual motion. Neuron, 73(6), 1228–1240. doi: 10.1016/j.neuron.2012.01.022

Fischer, E., Bulthoff, H. H., Logothetis, N. K., & Bartels, A. (2012b). Visual motion responses in the posterior cingulate sulcus: a comparison to V5/MT and MST. Cerebral cortex (New York, N.Y.: 1991), 22, 865–876. doi: 10.1093/cercor/bhr154

Fox, C. J., Iaria, G., & Barton, J. J. (2009). Defining the face processing network: optimization of the functional localizer in fMRI. Hum Brain Mapp, 30(5), 1637–1651. doi:http://dx.doi.org/10.1002/hbm.20630

Furlan, M., Wann, J. P., & Smith, A. T. (2014). A Representation of Changing Heading Direction in Human Cortical Areas pVIP and CSv. Cerebral Cortex, 24(11), 2848–2858. doi: 10.1093/cercor/bht132

Galletti, C., & Battaglini, P. P. (1989). Gaze-dependent visual neurons in area V3A of monkey prestriate cortex. J Neurosci, 9(4), 1112–1125.

Galletti, C., Battaglini, P. P., & Fattori, P. (1990). ‘Real-motion’ cells in area V3A of macaque visual cortex. Exp Brain Res, 82(1), 67–76.

Galletti, C., Battaglini, P. P., & Fattori, P. (1995). Eye position influence on the parieto-occipital area PO (V6) of the macaque monkey. Eur J Neurosci, 7(12), 2486–2501.

Galletti, C., & Fattori, P. (2003). Neuronal mechanisms for detection of motion in the field of view. Neuropsychologia, 41(13), 1717–1727.

Galletti, C., Fattori, P., Battaglini, P. P., Shipp, S., & Zeki, S. (1996). Functional demarcation of a border between areas V6 and V6A in the superior parietal gyrus of the macaque monkey. Eur J Neurosci, 8(1), 30–52.

Galletti, C., Fattori, P., Gamberini, M., & Kutz, D. F. (1999). The cortical visual area V6: brain location and visual topography. Eur J Neurosci, 11(11), 3922–3936.

Galletti, C., Gamberini, M., Kutz, D. F., Fattori, P., Luppino, G., & Matelli, M. (2001). The cortical connections of area V6: an occipito-parietal network processing visual information. Eur J Neurosci, 13(8), 1572–1588.

Ilg, U. J., Schumann, S., & Thier, P. (2004). Posterior parietal cortex neurons encode target motion in world-centered coordinates. Neuron, 43(1), 145–151. doi: 10.1016/j.neuron.2004.06.006

Ilg, U. J., & Thier, P. (2003). Visual tracking neurons in primate area MST are activated by smooth-pursuit eye movements of an “imaginary” target. J Neurophysiol, 90(3), 1489–1502.

Inaba, N., Miura, K., & Kawano, K. (2011). Direction and speed tuning to visual motion in cortical areas MT and MSTd during smooth pursuit eye movements. Journal of neurophysiology, 105(4), 1531–1545. doi: 10.1152/jn.00511.2010

Kawano, K., Shidara, M., Watanabe, Y., & Yamane, S. (1994). Neural activity in cortical area MST of alert monkey during ocular following responses. Journal of Neurophysiology, 71(6), 2305–2324.

Kayser, C., Kording, K. P., & Konig, P. (2004). Processing of complex stimuli and natural scenes in the visual cortex. Curr Opin Neurobiol, 14(4), 468–473.

Kleiner, M., Brainard, D. H., Pelli, D., Ingling, A., Murray, R. M., & Broussard, C. (2007). What’s new in Psychtoolbox-3. Perception, 36(14), 1.1–16.

Krekelberg, B., Boynton, G. M., & van Wezel, R. J. (2006). Adaptation: from single cells to BOLD signals. Trends Neurosci, 29(5), 250–256.

Krekelberg, B., & van Wezel, R. J. (2013). Neural mechanisms of speed perception: transparent motion. Journal of neurophysiology, 110, 2007–2018. doi: 10.1152/jn.00333.2013

Lee, B., Pesaran, B., & Andersen, R. A. (2007). Translation speed compensation in the dorsal aspect of the medial superior temporal area. The Journal of neuroscience: the official journal of the Society for Neuroscience, 27(10), 2582–2591. doi:10.1523/jneurosci.3416-06.2007

Lindner, A., Schwarz, U., & Ilg, U. J. (2001). Cancellation of self-induced retinal image motion during smooth pursuit eye movements. Vision research, 41(13), 1685–1694.

Lingnau, A., Wall, M. B., & Smith, A. T. (2009). Speed encoding in human visual cortex revealed by fMRI adaptation Hiroshi Ashida. 9, 1–14. doi: 10.1167/9.13.3.Introduction

Liu, J., & Newsome, W. T. (2003). Functional organization of speed tuned neurons in visual area MT. Journal of neurophysiology, 89, 246–256. doi: 10.1152/jn.00097.2002

Liu, J., & Newsome, W. T. (2005). Correlation between speed perception and neural activity in the middle temporal visual area. The Journal of neuroscience: the official journal of the Society for Neuroscience, 25, 711–722. doi: 10.1523/JNEUROSCI.4034-04.2005

Malonek, D., Tootell, R. B. H., & Grinvald, A. (1994). Optical Imaging Reveals the Functional Architecture Of Neurons Processing Shape and Motion In Owl Monkey Area MT. Proceedings of the Royal Society (London) B, 258(1352), 109–119.

MATLAB. (2010).version 7.10.0 Natick, Massachusetts: The MathWorks Inc.

Maunsell, J. H., & Van Essen, D. C. (1983a). The connections of the middle temporal area and their relationship to a cortical hierarchy in the macaque monkey. Journal of Neuroscience, 3(12), 2563–2586.

Maunsell, J. H., & Van Essen, D. C. (1983b). Functional properties of neurons in middle temporal visual area of the macaque monkey. II. Binocular interactions and sensitivity to binocular disparity. J Neurophysiol, 49(5), 1148–1167.

McKeefry, D. J., Burton, M. P., Vakrou, C., Barrett, B. T., & Morland, A. B. (2008). Induced deficits in speed perception by transcranial magnetic stimulation of human cortical areas V5/MT+ and V3A. J Neurosci, 28(27), 6848–6857. doi: 28/27/6848 [pii] 10.1523/JNEUROSCI.1287-08.2008

Mikami, A., Newsome, W. T., & Wurtz, R. H. (1986). Motion selectivity in macaque visual cortex I. Mechanisms of speed and directional selectivity in extrastriate area MT. Journal of Neurophysiology, 55, 1308–1327.

Murray, S. O., & Wojciulik, E. (2004). Attention increases neural selectivity in the human lateral occipital complex. Nat Neurosci, 7(1), 70–74. doi: http://dx.doi.org/10.1038/nn1161

Newsome, W. T., Mikami, A., & Wurtz, R. H. (1986). Motion Selectivity In Macaque Visual-Cortex .3. Psychophysics and Physiology Of Apparent Motion. Journal of Neurophysiology, 55(6), 1340–1351.

Newsome, W. T., Wurtz, R. H., Dürsteler, M. R., & Mikami, A. (1985). Deficits in visual motion processing following ibotenic acid lesions of the middle temporal visual area of the macaque monkey. Journal of Neuroscience, 5(3), 825–840.

Nover, H., Anderson, C.H., & DeAngelis, G. C. (2005). A logarithmic, scale-invariant representation of speed in macaque middle temporal area accounts for speed discrimination performance. (1529-2401 (Electronic)).

Olson, C. R., Musil, S. Y., & Goldberg, M. E. (1996). Single neurons in posterior cingulate cortex of behaving macaque: eye movement signals. J Neurophysiol, 76(5), 3285–3300.

Orban, G. A., de Wolf, J., & Maes, H. (1984). Factors influencing velocity coding in the human visual system. Vision research, 24(1), 33–39.

Orban, G. A., Lagae, L., Raiguel, S., Xiao, D., & Maes, H. (1995). The speed tuning of medial superior temporal (MST) cell responses to optic-flow components. Perception, 24(3), 269–285.

Orban, G. A., Saunders, R. C., & Vandenbussche, E. (1995). Lesions Of the Superior Temporal Cortical Motion Areas Impair Speed Discrimination In the Macaque Monkey. European Journal Of Neuroscience, 7(11), 2261–2276.

Pasternak, T., & Merigan, W. H. (1994). Motion perception following lesions of the superior tempora lsulcus in the monkey. Cerebral Cortex, 4, 247–259.

Perrone, J. A., & Thiele, A.. (2001). Speed skills: measuring the visual speed analyzing properties of primate MT neurons. Nat Neurosci, 4(5), 526–532.

Pitzalis, S., Galletti, C., Huang, R. S., Patria, F., Committeri, G., Galati, G., … Sereno, M. I. (2006). Wide-field retinotopy defines human cortical visual area v6. The Journal of neuroscience: the official journal of the Society for Neuroscience, 26, 7962–7973. doi: 10.1523/JNEUROSCI.0178-06.2006

Pitzalis, S., Sdoia, S., Bultrini, A., Committeri, G., Di Russo, F., Fattori, P., Galati, G. (2013). Selectivity to translational egomotion in human brain motion areas. PloS one, 8, e60241. doi: 10.1371/journal.pone.0060241

Pitzalis, S., Sereno, M. I., Committeri, G., Fattori, P., Galati, G., Patria, F., & Galletti, C. (2010). Human v6: the medial motion area. Cerebral cortex (New York, N.Y.: 1991), 20, 411–424. doi: 10.1093/cercor/bhp112

Pitzalis, S., Strappini, F., De Gasperis, M., Bultrini, A., & Di Russo, F. (2012). Spatio-temporal brain mapping of motion-onset VEPs combined with fMRI and retinotopic maps. PloS one, 7, e35771. doi: 10.1371/journal.pone.0035771

Priebe, N. J., Cassanello, C. R., & Lisberger, S. G. (2003). The Neural Representation of Speed in Macaque Area MT/V5. The Journal of neuroscience: the official journal of the Society for Neuroscience, 23(13), 5650–5661.

Priebe, N. J., & Lisberger, S. G. (2004). Estimating Target Speed from the Population Response in Visual Area MT. The Journal of neuroscience: the official journal of the Society for Neuroscience, 24(8), 1907–1916. doi: 10.1523/JNEUROSCI.4233-03.2004

Priebe, N. J., Lisberger, S. G., & Movshon, J. A. (2006). Tuning for spatiotemporal frequency and speed in directionally selective neurons of macaque striate cortex. The Journal of Neuroscience, 26(11), 2941–2950.

Quinlan, D. J., & Culham, J. C. (2007). fMRI reveals a preference for near viewing in the human parieto-occipital cortex. NeuroImage, 36(1), 167–187. doi: 10.1016/j.neuroimage.2007.02.029

Rodman, H. R., & Albright, T. D. (1987). Coding of visual stimulus velocity in area MT of the macaque. Vision Research, 27(12), 2035–2048.

Shenoy, K. V., Crowell, J. A., & Andersen, R. A. (2002). Pursuit speed compensation in cortical area MSTd. J Neurophysiol, 88(5), 2630–2647.

Singh, K. D., Smith, A. T., & Greenlee, M. W. (2000). Spatiotemporal frequency and direction sensitivities of human visual areas measured using fMRI. Neuroimage, 12(5), 550–564.

Smith, A. T., Wall, M. B., & Thilo, K. V. (2012). Vestibular inputs to human motion-sensitive visual cortex. Cerebral cortex (New York, N.Y.: 1991), 22, 1068–1077. doi: 10.1093/cercor/bhr179

Sunaert, S., Van Hecke, P., Marchal, G., & Orban, G. A. (2000). Attention to speed of motion, speed discrimination, and task difficulty: an fMRI study. Neuroimage, 11(6 Pt 1), 612–623.

Tanaka, K., & Saito, H. (1989). Analysis of motion of the visual field by direction, expansion/contraction, and rotation cells clustered in the dorsal part of the medial superior temporal area of the macaque monkey. Journal of Neurophysiology, 62(3), 626–641.

Tootell, R. B. H., Mendola, J. D., Hadjikhani, N. K., Ledden, P. J., Liu, A. K., Reppas, J. B., … Dale, A. M. (1997). Functional analysis of V3A and related areas in human visual cortex. J Neurosci, 17(18), 7060–7078.

Tootell, R. B. H., Reppas, J. B., Kwong, K. K., Malach, R., Born, R. T., Brady, T. J., … Belliveau, J. W. (1995). Functional analysis of human MT and related visual cortical areas using magnetic resonance imaging. Journal Of Neuroscience, 15(4), 3215–3230.

Ungerleider, L. G., & Desimone, R. (1986). Cortical connections of visual area MT in the macaque. Journal of Comparative Neurology, 248, 190–222.

Van Dijk, K. R., Hedden, T., Venkataraman, A., Evans, K. C., Lazar, S. W., & Buckner, R. L. (2010). Intrinsic functional connectivity as a tool for human connectomics: theory, properties, and optimization. J Neurophysiol, 103(1), 297–321. doi: http://dx.doi.org/10.1152/jn.00783.2009

Vinje, W. E., & Gallant, J. L. (2000). Sparse coding and decorrelation in primary visual cortex during natural vision. Science, 287(5456), 1273–1276.

Wall, M. B., & Smith, A.T. (2008). The representation of egomotion in the human brain. Current biology: CB, 18, 191–194. doi: 10.1016/j.cub.2007.12.053

Yamasaki, D. S., & Wurtz, R. W. (1991). Recovery of function after lesions in the superior temporal sulcus in the monkey. Journal of Neurophysiology, 66(3), 651–673.

Zhang, T., Heuer, H. W., & Britten, K. H. (2004). Parietal area VIP neuronal responses to heading stimuli are encoded in head-centered coordinates. Neuron, 42(6), 993–1001.

